# A highly attenuated vaccinia virus strain LC16m8-based vaccine for severe fever with thrombocytopenia syndrome

**DOI:** 10.1101/2020.08.06.239368

**Authors:** Tomoki Yoshikawa, Satoshi Taniguchi, Hirofumi Kato, Naoko Iwata-Yoshikawa, Hideki Tani, Takeshi Kurosu, Hikaru Fujii, Natsumi Omura, Miho Shibamura, Shumpei Watanabe, Kazutaka Egawa, Takuya Inagaki, Satoko Sugimoto, Supranee Phanthanawiboon, Shizuko Harada, Souichi Yamada, Shuetsu Fukushi, Shigeru Morikawa, Noriyo Nagata, Masayuki Shimojima, Masayuki Saijo

**Affiliations:** Department of Virology 1, National Institute of Infectious Diseases, Musashimurayama-shi, Tokyo, Japan; Department of Pathology, National Institute of Infectious Diseases, Musashimurayama-shi, Tokyo, Japan; Department of Virology, Toyama Institute of Health, Imizu-shi, Toyama, Japan; Department of Microbiology, Faculty of Veterinary Medicine, Okayama University of Science, Imabari-shi, Ehime, Japan

## Abstract

Severe fever with thrombocytopenia syndrome (SFTS) caused by Dabie bandavirus (formerly SFTS virus [SFTSV]) is an emerging hemorrhagic infectious disease with a high case-fatality rate. One of the best strategies for preventing SFTS is to develop a vaccine, which is expected to induce both humoral and cellular immunity. We applied a highly attenuated but still immunogenic vaccinia virus strain LC16m8 (m8) as a recombinant vaccine for SFTS. Recombinant m8s expressing SFTSV nucleoprotein (m8-N), envelope glycoprotein precursor (m8-GPC), and both N and GPC (m8-N+GPC) in the infected cells were generated. Both m8-GPC- and m8-N+GPC-infected cells were confirmed to produce SFTSV-like-particles (VLP) *in vitro*, and the N was incorporated in the VLP produced by the infection of cells with m8-N+GPC. Specific antibodies to SFTSV were induced in mice inoculated with each of the recombinant m8s, and the mice were fully protected from lethal challenge with SFTSV at both 10^3^ TCID_50_ and 10^5^ TCID_50_. In mice that had been immunized with vaccinia virus strain Lister in advance of m8-based SFTSV vaccine inoculation, protective immunity against the SFTSV challenge was also conferred. The pathological analysis revealed that mice immunized with m8-GPC or m8-N+GPC did not show any histopathological changes without any viral antigen-positive cells, whereas the control mice showed focal necrosis with inflammatory infiltration with SFTSV antigen-positive cells in tissues after SFTSV challenge. The passive serum transfer experiments revealed that sera collected from mice inoculated with m8-GPC or m8-N+GPC but not with m8-N conferred protective immunity against lethal SFTSV challenge in naïve mice. On the other hand, the depletion of CD8-positive cells *in vivo* did not abrogate the protective immunity conferred by m8-based SFTSV vaccines. Based on these results, the recombinant m8-GPC and m8-N+GPC were considered promising vaccine candidates for SFTS.

**Author Summary:** Severe fever with thrombocytopenia syndrome (SFTS) is an emerging viral hemorrhagic fever with a high case-fatality rate (approximately 5% to >40%). Indigenous SFTS has been reported in China, Japan, South Korea, and Vietnam. Thus, the development of an effective vaccine for SFTS is urgently needed. Vaccinia virus (VAC) was previously used as a vaccine for smallpox. Unfortunately, after these strains, the so-called second generation of VAC used during the eradication campaign was associated with severe adverse events, and the third generation of VAC strains such as LC16m8 (m8) and modified vaccinia Ankara (MVA) was established. m8 is confirmed to be highly attenuated while still maintaining immunogenicity. m8 is licensed for use in healthy people in Japan. At the present time, approximately 100,000 people have undergone vaccination with m8 without experiencing any severe postvaccine complications. At present, third-generation VAC strains are attractive for a recombinant vaccine vector, especially for viral hemorrhagic infectious diseases, such as Ebola virus disease, Lassa fever, Crimean-Congo hemorrhagic fever, and SFTS. We investigated the practicality of an m8-based recombinant vaccine for SFTS as well as other promising recombinant VAC-based vaccines for viral hemorrhagic infectious diseases.

## Introduction

Severe fever with thrombocytopenia syndrome (SFTS) is an emerging viral hemorrhagic fever with a high case-fatality rate (approximately 5 to over 40%) [1–6]. The clinical symptoms are in general fever, malaise, myalgia, nausea, vomiting, and diarrhea. The laboratory findings include leukocytopenia and thrombocytopenia in the total blood cell counts, and elevated serum levels of hepatic enzymes [1, 7–11]. The disease is caused by Dabie bandavirus, which had been called SFTS virus (SFTSV), a novel tick-borne virus in the order *Bunyavirales*, family *Phenuiviridae*, and genus *Bandavirus*. The viral genome consists of three negative-stranded RNA segments and encodes four genes: RNA-dependent RNA polymerase, envelope glycoproteins precursor (GPC), nucleoprotein (N), and nonstructural protein. Indigenous SFTS had been reported in China, Japan, South Korea, and Vietnam [1–3, 12–14]. Hence the development of an effective vaccine for SFTS is urgently needed. Thus far, it has been reported that a live recombinant vesicular stomatitis virus or a DNA vaccine expressing the SFTSV glycoproteins (GPs) originated from SFTSV GPC gene-elicited protective immunity against SFTSV in lethal mouse or ferret models [15, 16].

Vaccinia virus (VAC) was used as the smallpox vaccine, and smallpox vaccines produced by using a variety of VAC strains were used during the global smallpox eradication program led by the World Health Organization. Since smallpox eradication was declared in 1980, VAC has been used as a recombinant vaccine vector with an expectation of immunogenicity [17, 18]. However, since the VAC strains used as the vaccine vector in the beginning of the eradication campaign were mainly so-called second-generation smallpox vaccines that were associated with severe side effects, such as encephalitis, encephalopathy, conjunctivitis, progressive vaccinia, eczema vaccinatum and generalized fetal VAC infections [17, 19, 20], the application of VAC has not been popular. The VAC strain LC16m8 (m8), which is categorized as a third-generation smallpox vaccine, as well as the modified vaccinia Ankara (MVA), is confirmed to have a highly attenuated phenotype but to maintain immunogenicity to protect against other orthopoxvirus infections, such as monkeypox [21–23]. m8 is licensed for use in healthy people in Japan, and approximately 100,000 people have been vaccinated with m8 thus far, with antibody response comparable to the first generation vaccine [24] and without experiencing severe postvaccine complications [24, 25]. This well-balanced safety and immunogenicity profile of m8 represents a significant advantage over the first- and second generations of VAC. In the present study, recombinant m8 strains that express SFTSV N (m8-N), GPC (m8-GPC), or both N and GPC (m8-N+GPC) (m8-based SFTSV vaccines) were generated as SFTS vaccine candidates. It is hypothesized that N and GPC contributed to cellular immunity and/or humoral immunity against SFTSV infection. The immunogenicity and protective efficacy of m8-based SFTSV vaccines against SFTSV were evaluated using a mouse model of lethal SFTS.

## Results

### Generation and characterization of recombinant m8 harboring SFTSV genes

The final recombinant m8-based SFTSV vaccines harbored the SFTSV gene expression cassette in the flanking region between the B4R and B6R genes in place of the B5R gene (Fig. 1A). The SFTSV gene expression in RK13 cells infected with m8-N, m8-GPC, or m8-N+GPC was confirmed by an indirect immunofluorescence assay (IFA) (Fig. 1B). VLP production in the m8-GPC or m8-N+GPC-infected cells was evaluated by electron microscopy (Fig. 1C). The morphological characteristics of VLP were enveloped and spherical with approximate diameters of 100 nm and seemed similar to that of SFTSV. Next, the incorporation of the N protein into the VLP, which was produced from cells infected with m8-N+GPC, was evaluated (Fig. 1D). When the VLP was pulled down in immunoprecipitation using anti-SFTSV Gc antibodies, the amount of N protein recovered from cells infected with m8-N+GPC was more extensive than that of m8-N-infected cells in comparison to when no antibodies were used. On the other hand, the addition of the anti-SFTSV Gc antibody did not affect the recovery of the N protein from the m8-N-infected-cell supernatant, which did not contain the VLP. This result indicated that the m8-GPC- or m8-N+GPC-infected cells produced SFTSV-like particles, and that the N protein was incorporated into the VLP when cells were infected with m8-N+GPC.

**Fig 1.**
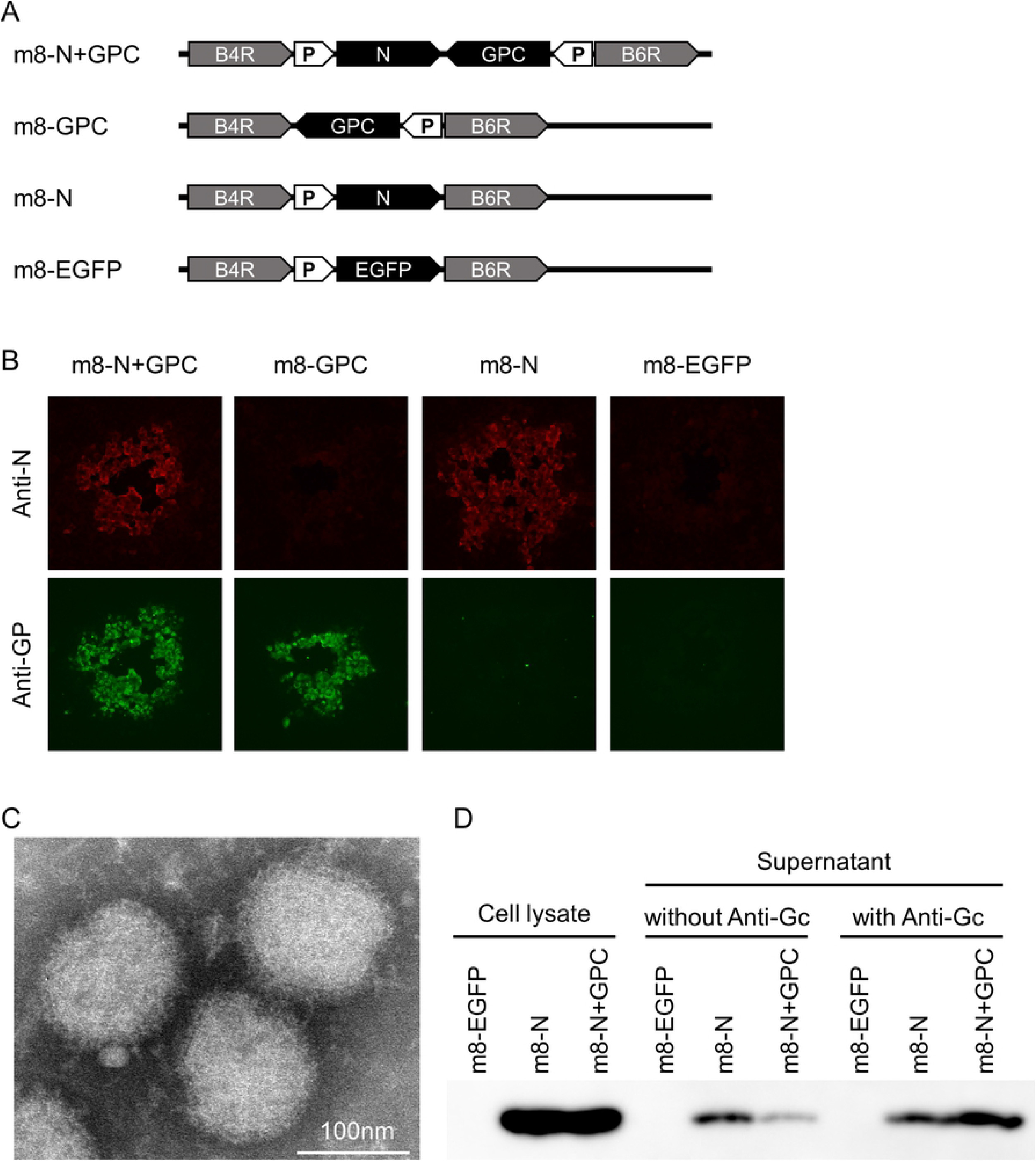
The construction, generation, and characterization of recombinant LC16m8 harboring SFTSV N, GPC, and both N and GPC, and EGFP genes. A schematic illustration of the site harboring EGFP, SFTSV genes in the genome in m8-EGFP, or m8-based SFTSV vaccines (A). The VAC B5R gene, flanked by the B4R and B6R genes, was substituted for the SFTSV gene. Synthetic vaccinia early/late promoter is indicated as P in the arrow-shaped box. The expression of SFTSV N protein and GPs was confirmed in the cells infected with m8-based SFTSV vaccines (B). Purified VLP from the supernatant of the m8-N+GPC-infected cell was negatively stained and observed under a transmission electron microscope (C). The supernatant was precipitated using PEG 6000, and the VLP was banded by ultracentrifugation at the interface of 20% and 60% (w/v) double-cushion sucrose. RK13 cells were infected with m8-N+GPC at an MOI of 0.1 per cell, and the culture supernatant was collected at 3 DPI. The collected VLP in the culture supernatant was banded at the 20/60% interface and then negatively stained with 2% phosphotungstic acid. The bar in the image indicates the length of 100 nm. The picture of the VLP was derived from the supernatant of the m8-N+GPC-infected cells. The morphology of SFTS VLP from the m8-GPC-infected cells was indistinguishable from that of m8-N+GPC. SFTS N protein incorporation into the VLP was evaluated (D). The VLP in the supernatant from the RK13 cells infected with m8-EGFP, m8-N, or m8-N+GPC was reacted with or without anti-SFTSV GPC mAb (clone C6C1) and then precipitated with anti-mouse IgG magnetic beads. The precipitated magnetic beads were then resolved on SDS-PAGE and analyzed by Western blotting using rabbit anti-SFTSV N polyclonal antibody. RK13 cell lysates infected with m8-EGFP, m8-N, or m8-N+GPC were shown as a positive control.

### Immunogenicity of m8-based SFTSV vaccines in mice

Groups (5 per group) of seven- to 8-week-old naïve Type I interferon alpha receptor-deficient (IFNAR-/-) mice were subcutaneously inoculated twice at a 2-week interval with either m8-based SFTSV vaccine or m8-EGFP as a negative control. None of the mice vaccinated with any of the m8-based SFTSV vaccines, including the negative control, developed observable clinical signs of illness, such as ruffled fur or pock formation at the site of inoculation. To evaluate the immunogenicity of m8-based SFTSV vaccines, sera were collected from IFNAR-/- mice and subjected to an IFA and N.T. assay. Five naïve IFNAR-/- mice were subcutaneously infected with 10 TCID_50_ of SFTSV YG-1 as a positive control. Since one of the five mice died at 13 days post-SFTSV infection, sera were collected from four mice at four weeks after infection. In addition, one of the sera collected from four SFTSV-infected mice was omitted from the subsequent analyses, since the serum did not seroconvert (i.e., both IFA and N.T. results were negative). The specific IgG against SFTSV N or GPs in the sera were elicited by the infection of mice with m8-N, m8-GPC, or m8-N+GPC (Fig. 2A, B). The N.T. antibodies to SFTSV were also elicited, and the 50% N.T. antibody titers of the sera collected from m8-GPC-infected, m8-N+GPC-infected, and even SFTSV-infected mice were approximately 1:10 dilution (Fig. 2C). These results indicated that m8-based SFTSV vaccines induced humoral immunity, and the induction potency was comparable to that induced by the authentic SFTSV.

**Fig 2.**
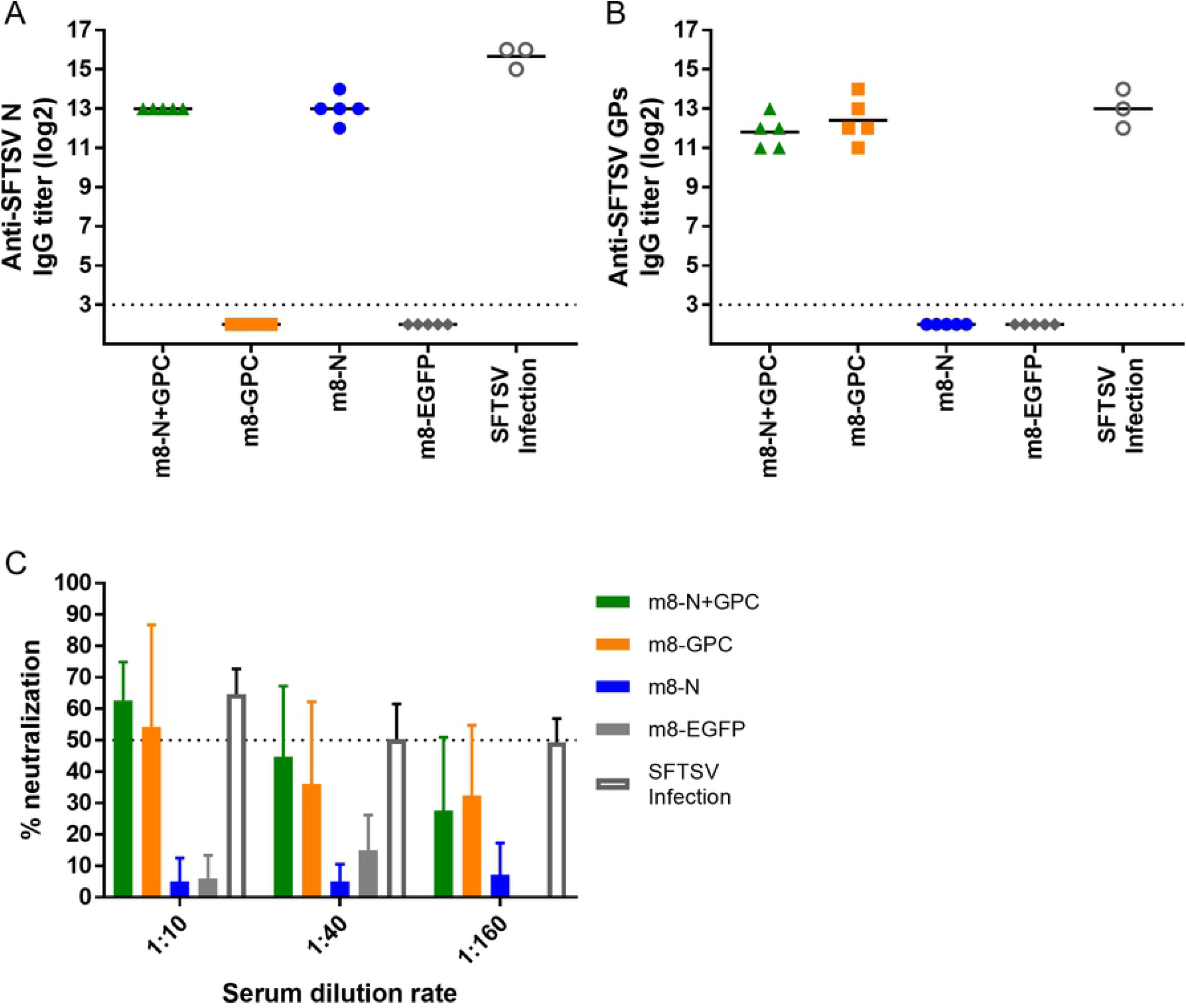
Immunogenicity of the m8-based SFTSV vaccines in mice. Naïve IFNAR-/- mice were subcutaneously inoculated twice at a 2-week interval with a dose of 1 × 10^6^ PFU of m8-EGFP, m8-N, m8-GPC or m8-N+GPC or subcutaneously infected with 10 TCID_50_ of SFTSV YG-1 (SFTSV infection). Two weeks after the second inoculation of mice with the recombinant m8s or 4 weeks after SFTSV infection, the mice were euthanized and sacrificed. Blood was collected to evaluate Anti-SFTSV N-specific (A) or GPs-specific (B) IgG production. Each dot represents the antibody titer. Horizontal lines represent the mean value. The horizontal dotted line at “2^3^” indicates the detection limit. The neutralizing antibodies to SFTSV were also measured (C). SFTSV YG-1 was mixed and incubated with 3 serial 4-fold dilutions (1/10, 1/40, or 1/160) of serum. The percent neutralization titer was determined to divide the number of foci by that of naïve mice.

### Vaccine efficacy of m8-based SFTSV vaccines in mice

Groups (8 to 10 per group) of six- to nine-week-old IFNAR-/- mice were subcutaneously inoculated twice at a two-week interval with each m8-based SFTSV vaccine. At two weeks after the second inoculation, the mice were subcutaneously challenged with 1 × 10^3^ or 1 × 10^5^ TCID_50_ of SFTSV YG-1. While almost all of the control mice died after the development of clinical signs, such as ruffled fur, hunched posture, and weight loss, all of the mice inoculated with either m8-GPC or m8-N+GPC survived without developing any obvious clinical signs during the two-week observation period after the SFTSV challenge (Fig. 3A-D). Furthermore, all the mice that had been inoculated with m8-N survived without developing any obvious clinical signs after SFTSV challenge with a dose of 1 × 10^3^ TCID_50_ during the two-week observation period (Fig. 3A, C). On the other hand, the mice inoculated with m8-N survived but transiently developed clinical signs, when challenged with 1 × 10^5^ TCID_50_ of SFTSV (Fig. 3B, D). To assess the immune response to N and GPs of SFTSV, sera collected from mice that survived two weeks after the SFTSV infection were subjected to an IFA to measure specific IgG against SFTSV N and GPs, which would not be induced by monovalent m8-GPC and m8-N, respectively. SFTSV GPs-specific IgG was not induced in the m8-N inoculated mice, while N-specific IgG was induced in five of eight m8-GPC-inoculated mice when challenged with 1 × 10^3^ TCID_50_ SFTSV (Fig. 3E). On the other hand, all mice inoculated with m8-N or m8-GPC seroconverted after infection with 1 × 10^5^ TCID_50_ of SFTSV (Fig. 3F). These results indicated that the m8-based SFTSV vaccines conferred protection against a lethal SFTSV challenge. There was a difference in the degree of the efficacy between m8-N and m8-GPC including m8-N+GPC.

**Fig 3.**
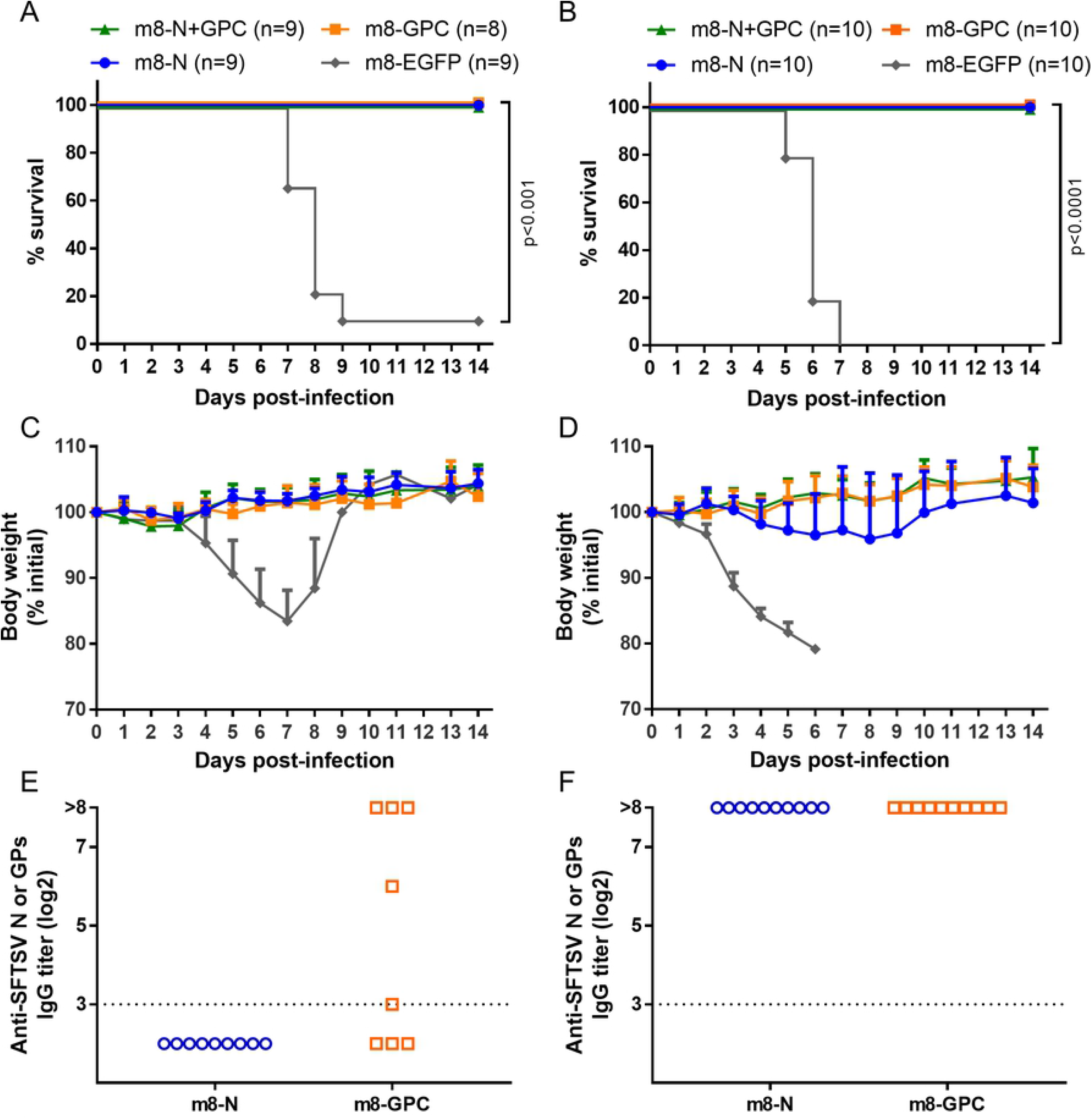
Survival, weight change, and seropositivity in m8-based SFTSV vaccine-inoculated mice followed by lethal SFTSV challenge. IFNAR-/- mice were subcutaneously inoculated twice at a 2-week interval with a dose of 1 × 10^6^ PFU of m8-EGFP, m8-N, m8-GPC or m8-N+GPC. Two weeks later, from the second inoculation of mice with m8-EGFP or m8-based SFTSV vaccines, the mice were subcutaneously challenged with 1 × 10^3^ (A, C) or 1 × 10^5^ (B, D) TCID_50_ of SFTSV YG-1. Survival (A, B) and the sequential change in percent weight from the initial (C, D) were followed daily for 2 weeks. The number of mice in a group is shown as “n” in the legend. The log-rank test was used to determine the level of statistical significance. The calculated p-values are shown beside the groups that were compared. Seropositivity in mice inoculated with m8-N or m8-GPC then infected with SFTSV was evaluated. Serum was collected from the m8-N and m8-GPC-inoculated mice at 2 weeks after 1 x 10^3^ (E) or 1 × 10^5^ (F) TCID_50_ of SFTSV infection. Specific IgG against SFTSV GPs or N, which should be induced by SFTSV infect, and which should never be induced by monovalent m8-N or m8-GPC, was measured. Each dot represents the antibody titer. The horizontal dotted line at 2^3^ indicates the detection limit.

### Effect of preexisting immunity against VAC

To evaluate the impact of preexisting immunity to VAC on the protective immunity induced by m8-based SFTSV vaccines, IFNAR-/- mice were immunized with VAC and then inoculated with each m8-based SFTSV vaccine and challenged with SFTSV. The survival rate of the VAC-preimmunized mice infected with 1 × 10^3^ or 1 × 10^5^ TCID_50_ of SFTSV was improved by inoculation with m8-N, m8-GPC, or m8-N+GPC in comparison to control mice inoculated with m8-EGFP (Fig. 4). However, the survival rates were lower than those of the mice that were not immunized with VAC Lister in advance of the vaccinations (Fig. 3 A, B, and Fig. 4A, B). The survival rate of m8-N+GPC-inoculated mice was similar to that of m8-GPC-inoculated mice, whereas the survival rate of m8-N-inoculated mice was lower in comparison to m8-N+GPC- and m8-GPC-inoculated mice. The disease severity of the groups tended to show a similar trend to the survival rate. Although all the mice, even those inoculated with m8-N, m8-GPC, m8-N+GPC in advance, developed clinical symptoms after infection with 1 × 10^3^ TCID_50_ or 1 × 10^5^ TCID_50_ of SFTSV, the weight loss in the mice inoculated with m8-based SFTSV vaccines, especially those inoculated with m8-GPC or m8-N+GPC, was milder in comparison to mice inoculated with m8-EGFP (Fig. 4C, D). These results indicated that m8-based SFTSV vaccines conferred protection against lethal SFTSV challenge in mice, even when preexisting immunity to VAC was present.

**Fig 4.**
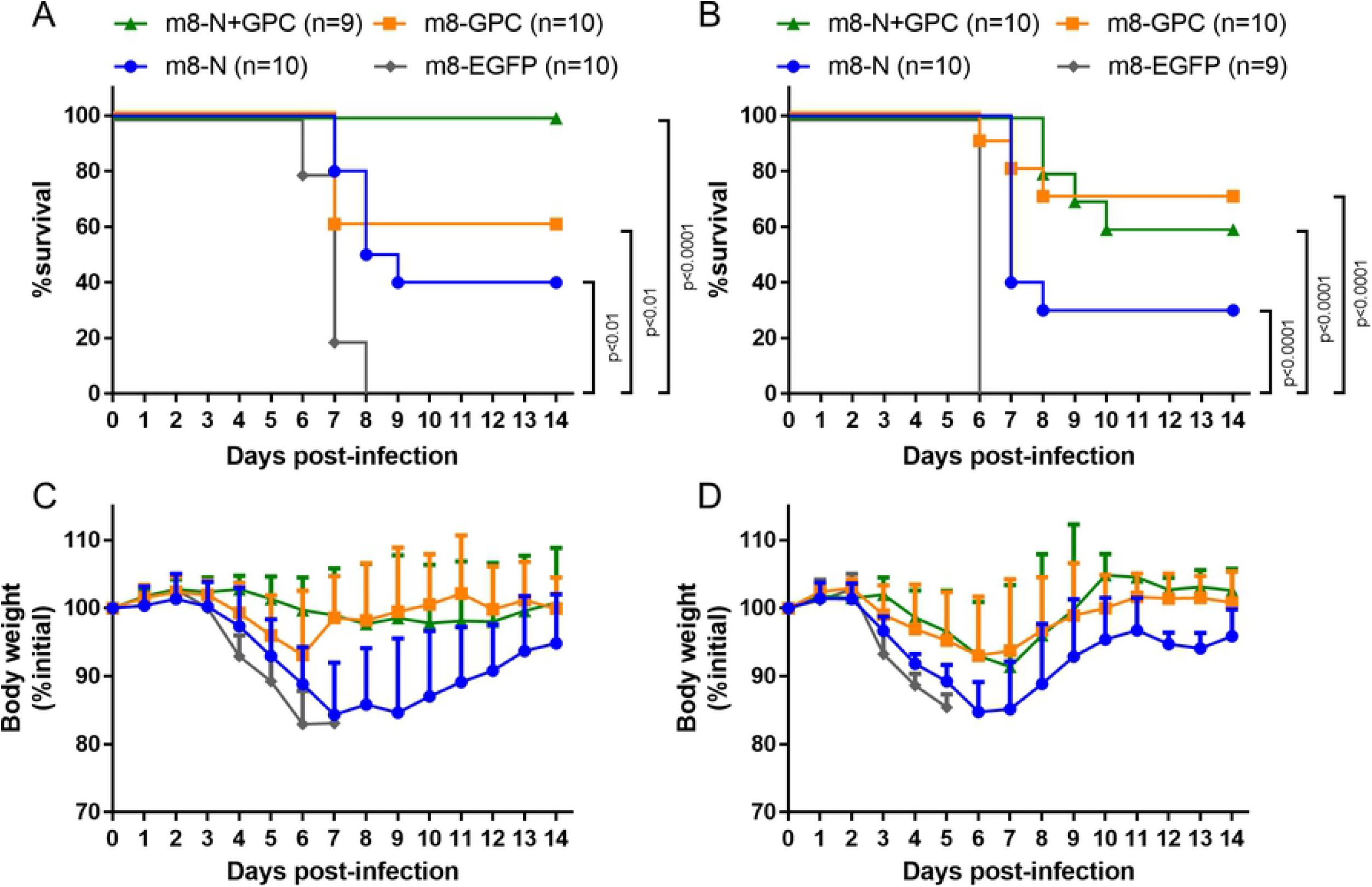
Survival and weight change in mice immunized with VAC before m8-based SFTSV vaccine inoculation. IFNAR-/- mice were inoculated twice at a 2-week interval with a dose of 1 x 10^6^ PFU of m8-EGFP, m8-N, m8-GPC or m8-N+GPC, 1 month after inoculation with 1 × 10^6^ PFU VAC strain Lister. Two weeks after the second inoculation of m8-EGFP, m8-N, m8-GPC, or m8-N+GPC, the mice were subcutaneously challenged with 1 × 10^3^ (A, C) or 1 × 10^5^ (B, D) TCID_50_ of SFTSV YG-1. Survival (A, B) and percent weight sequential change from the initial (C, D) were followed daily for 2 weeks. The number of mice used in a group is shown as “n” in the legend. The log-rank test with Bonferroni correction was used to determine the level of statistical significance. To adjust the significance level, the basic significance level was divided by the number of null hypotheses tested (i.e., 0.05/3=0.017). The calculated p-values are shown beside the groups that were compared.

### SFTSV infection dynamics in mice inoculated with m8-based SFTSV vaccines

To investigate the effect of m8-based SFTSV vaccines in mice on spatial and temporal SFTSV infection dynamics, 6-week-old IFNAR-/- mice (three per group on each collection day) were subcutaneously inoculated twice at a two-week interval with m8-based SFTSV vaccines. Two weeks after the second inoculation, the mice were subcutaneously challenged with 1 × 10^5^ TCID_50_ of SFTSV YG-1, and the sera and tissues were collected at 1, 3, and 5 DPI. Sera collected from mice inoculated with m8-EGFP showed a high SFTSV titer (Fig. 5A). In contrast, in sera from the mice inoculated with m8-N, m8-GPC, or m8-N+GPC, SFTSV was below the limit of detection or a tiny amount of the virus was detected (Fig. 5A). SFTSV N gene copies were detected in the sera in one-third of the m8-EGFP inoculated mice at 1 DPI, and the copy number was drastically increased at 3 and 5 DPI (Fig. 5B). The average N gene copies in 1 ml of serum were 1 × 10^6.3^ and 1 ×10^7.5^ copies at 3 and 5 DPI, respectively. On the other hand, the N gene copy numbers in sera of mice inoculated with m8-based SFTSV vaccines contained a significantly lower in comparison to mice inoculated with m8-EGFP. However, the copy number transition differed among mice inoculated with m8-N, m8-GPC, and m8-N+GPC (Fig. 5B). The SFTSV N genes were detected in m8-N-inoculated mice over 3-5 days, although the number of copies was small. In the sera of mice inoculated with m8-GPC, the N gene had been detected at 3 DPI but was below the limit of detection at 5 DPI. The N gene copies were below the detection limit in the sera of the m8-N+GPC-inoculated mice not only at 3 DPI but also at 5 DPI.

**Fig 5.**
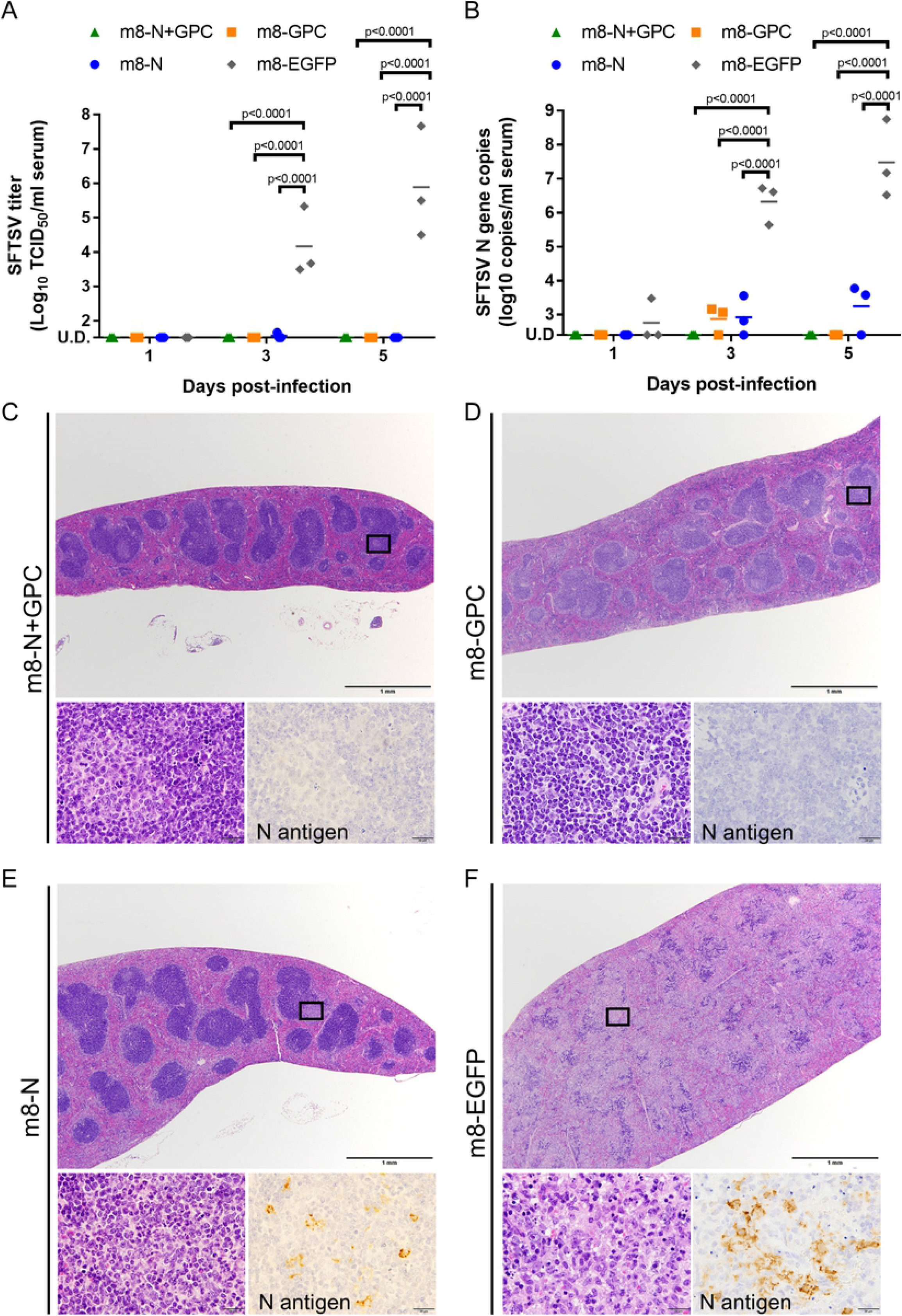
Infectious SFTSV and SFTSV genome levels and the histopathological changes in the tissue of m8-based vaccine-inoculated mice. IFNAR-/- mice were inoculated with m8-N, m8-GPC, m8-N+GPC, or m8-EGFP. At two weeks after the second inoculation, the mice were subcutaneously challenged with 1 × 10^5^ TCID_50_ of SFTSV YG-1. Three infected mice were sacrificed, and blood was collected at 1, 3, and 5 DPI. The sequential change in the infectious SFTSV titers (A) and SFTSV N gene copies (B) in the sera were measured by a standard TCID_50_ assay and qPCR. A two-way ANOVA with Sidak’s multiple-comparison test was used to determine the level of statistical significance. The calculated p-values are shown above the groups that were compared. U.D., under the detection limit, which is 32 TCID_50_ per ml or 125 copies per ml of serum. Histopathological changes in the spleen at 5 days of mice inoculated with m8-N+GPC (C), m8-GPC (D), m8-N (E), and m8-EGFP (F) after SFTSV challenge. High-magnification views of hematoxylin and eosin-stained tissue sections (left) and immunohistochemistry (right) of the areas of interest are shown at the bottom of the low magnification views of each group.

### Histopathological and immunohistochemical analysis

The liver, kidney, spleen, and cervical lymph nodes excised from mice inoculated with m8-N, m8-GPC, m8-N+GPC, and m8-EGFP after SFTSV challenge at 1, 3, and 5 DPI were subjected to histopathological and immunohistochemical analyses. The pathological changes of these organs differed among the 3 m8-based SFTSV vaccination groups and the control mice inoculated with m8-EGFP. There were no histopathological changes and any cells with viral antigen in the examined organs excised from m8-GPC or m8-N+GPC-inoculated mice (Table 1, 2, Fig. 5C and D). The m8-N-inoculated mice showed focal infiltration of lymphocytes in the liver at 1 and 3 DPI, although viral antigen-positive cells were not detected at that time. At 5 DPI, there were pathological changes, including multifocal necrosis in the liver, mild interstitial lymphocytic infiltration in the kidney, focal infiltration of neutrophils in the white pulp of the spleen, and depletion of lymphocytes in the cervical lymph nodes (Fig. 5E top and bottom left). Furthermore, in the m8-N-inoculated mice viral antigen-positive large mononuclear cells were detected only spleen and cervical lymph nodes, as was also observed in m8-EGFP-inoculated mice (Table 2 and Fig. 5E bottom right). In the control mice, the m8-EGFP-inoculated mice, the following histopathological changes were observed. Focal necrosis with infiltration of neutrophils and lymphocytes and mild infiltration of mononuclear cells were detected in the liver and kidney at 1 DPI. These hepatic and renal lesions gradually spread over time (Table 1). Lymphocyte depletion with apoptosis and infiltration of neutrophils were also detected in some white pulps in spleen and cervical lymph nodes at 3 DPI, and the lesions progressed as severe lymphocyte depletion with massive necrosis in the splenic white pulps, and cervical lymph nodes caused follicular structure almost disappeared on 5 DPI (Fig. 5F top and bottom left). Some viral antigen-positive cells that were large mononuclear cells began to be detected from 3DPI (Table 2 and Fig. 5F bottom right), and the cells were consistently localized with the lesions, including the liver, kidney, spleen, and lymph nodes. These results indicated that vaccination with either of the m8-based SFTSV vaccines conferred protective immunity that suppressed the propagation of SFTSV and inflammation *in vivo*. However, it is noteworthy that the efficacy induced by the m8-based SFTSV GPC vaccine is superior to that induced by vaccination with m8-N.

**Table 1.** Temporal change of lesion-positive tissues in SFTSV-challenged mice.

|  | Lesion (no. of positive / no. of mice) |  |  |  |  |  |  |  |  |  |  |  |
| --- | --- | --- | --- | --- | --- | --- | --- | --- | --- | --- | --- | --- |
| Tissue | Liver |  |  | Spleen |  |  | Kidney |  |  | Cervical LN |  |  |
| DPI | 1 | 3 | 5 | 1 | 3 | 5 | 1 | 3 | 5 | 1 | 3 | 5 |
| m8-EGFP | 3/3 | 3/3 | 3/3 | 0/3 | 2/3 | 3/3 | 3/3 | 3/3 | 3/3 | 0/3 | 1/3 | 3/3 |
| m8-N | 3/3 | 3/3 | 3/3 | 0/3 | 0/3 | 1/3 | 0/3 | 0/3 | 1/3 | 0/3 | 0/3 | 3/3 |
| m8-GPC | 0/3 | 0/3 | 0/3 | 0/3 | 0/3 | 0/3 | 0/3 | 0/3 | 0/3 | 0/3 | 0/3 | 0/3 |
| m8-G+N | 0/3 | 0/3 | 0/3 | 0/3 | 0/3 | 0/3 | 0/3 | 0/3 | 0/3 | 0/3 | 0/3 | 0/3 |

**Table 2.** Temporal change of SFTSV N antigen-positive tissues in SFTSV-challenged mice.

|  | SFTSV N Antigen (no. of positive / no. of mice) |  |  |  |  |  |  |  |  |  |  |  |
| --- | --- | --- | --- | --- | --- | --- | --- | --- | --- | --- | --- | --- |
| Tissue | Liver |  |  | Spleen |  |  | Kidney |  |  | Cervical LN |  |  |
| DPI | 1 | 3 | 5 | 1 | 3 | 5 | 1 | 3 | 5 | 1 | 3 | 5 |
| m8-EGFP | 0/3 | 2/3 | 3/3 | 0/3 | 3/3 | 3/3 | 0/3 | 1/3 | 3/3 | 0/3 | 2/3 | 3/3 |
| m8-N | 0/3 | 0/3 | 0/3 | 0/3 | 0/3 | 2/3 | 0/3 | 0/3 | 0/3 | 0/3 | 0/3 | 3/3 |
| m8-GPC | 0/3 | 0/3 | 0/3 | 0/3 | 0/3 | 0/3 | 0/3 | 0/3 | 0/3 | 0/3 | 0/3 | 0/3 |
| m8-G+N | 0/3 | 0/3 | 0/3 | 0/3 | 0/3 | 0/3 | 0/3 | 0/3 | 0/3 | 0/3 | 0/3 | 0/3 |

### Contribution of antibodies induced by m8-based SFTSV vaccine

To verify the contribution of the humoral immunity induced by m8-based SFTSV vaccines, sera obtained from the mice inoculated with each vaccine was passively transferred to naïve mice and challenged with SFTSV. As a control, a group of naïve mice was inoculated with sera obtained from mice three weeks after the infection with 1 × 10^1^ TCID_50_ of SFTSV. Six- to 10-week-old naïve IFNAR-/- mice (5 or 2 per group) underwent the intraperitoneal administration of sera, in which the presence of specific SFTSV antibodies was confirmed in the previous experiment (Fig. 2); then, they were immediately subcutaneously infected with SFTSV at a dose of 1 × 10^3^ TCID_50_. The survival rate mice treated with sera collected from mice immunized with each m8-based SFTSV vaccines showed no significant improvement in comparison to m8-EGFP inoculated mice (Fig. 6). However, there were one or two mice that did not develop any weight loss in the groups of mice treated with sera collected from m8-GPC-, m8-N+GPC-, or YG-1-inoculated mice. These sera contained specific and neutralizing antibodies against SFTSV (Fig. 2). These results suggested that the humoral immunity against SFTSV GPs contributed—to a certain degree—to the conferred anti-SFTSV protective immunity.

**Fig 6.**
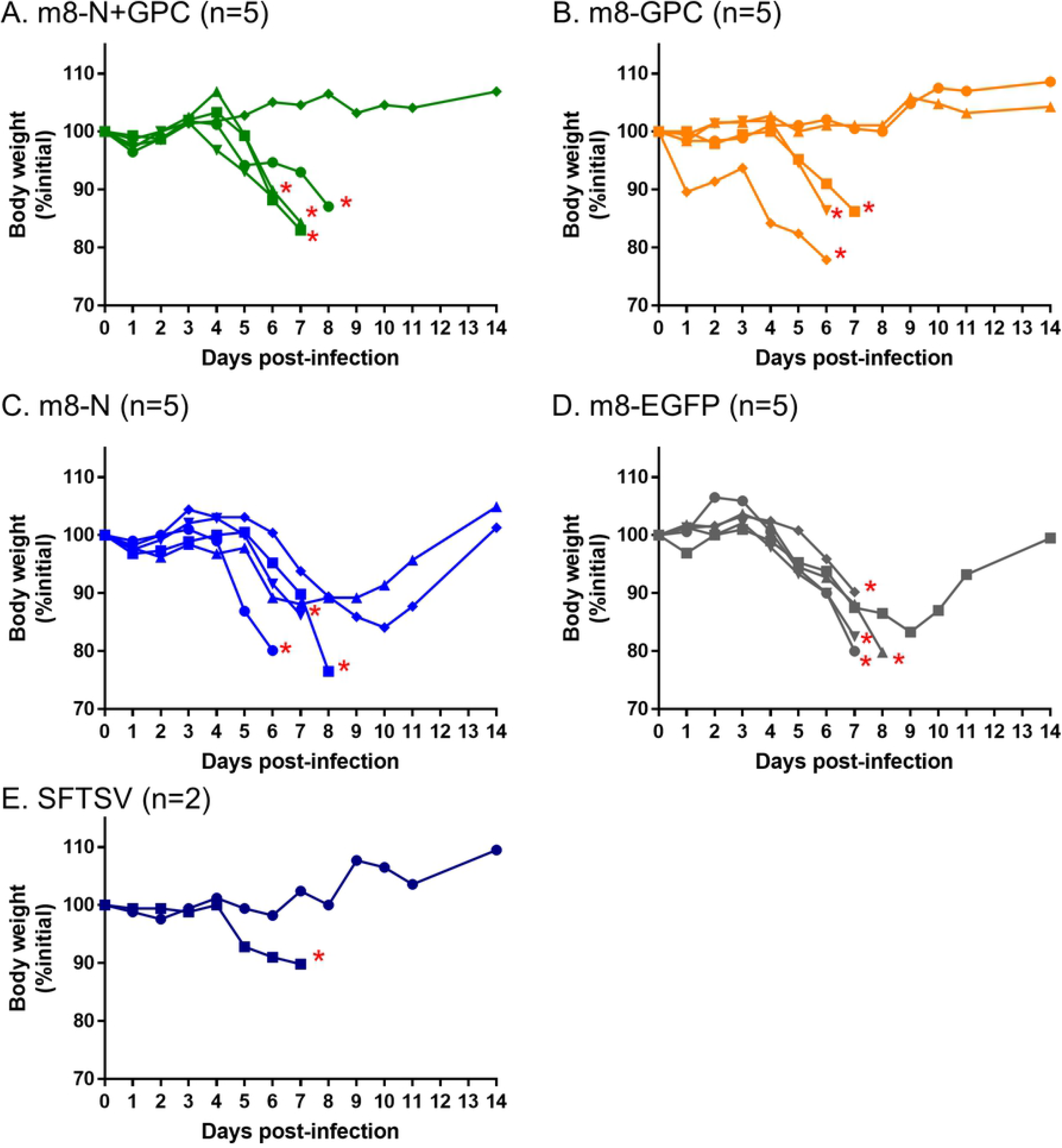
Weight change in mice to which sera collected from mice inoculated with each m8-based SFTSV vaccine was passively administered. IFNAR-/- mice were passively transferred with 100 μl of sera collected from mice inoculated m8-EGFP (A), m8-N (B), m8-GPC (C) or m8-N+GPC (D) at 1 day before and immediately before the subcutaneous challenge with SFTSV YG-1 at a dose of 1 × 10^3^ TCID_50_. As a positive control, sera obtained from mice 3 weeks after the infection with 10 TCID_50_ of SFTSV YG-1 was administered to a group of naïve mice (E). The percent weight change of each individual in the groups from the initial weight was measured daily for 2 weeks. The number of mice used in a group is shown as “n” in the legend. The asterisk indicates the endpoint of the individuals.

### Contribution of CD8-positive cells against SFTSV infections

CD8-positive cells were depleted *in vivo* during SFTSV challenge, which was performed to verify the contribution of CD8-positive cells, which are known to play a significant role in cellular immunity. The depletion of CD8-positive cells during the SFTSV challenge did not alter the survival rate in mice inoculated with either of the m8-based SFTSV vaccines or in the mice inoculated with the control m8-EGFP (Fig. 7). Furthermore, there was no marked difference in the sequential change in weight in the groups treated with anti-CD8 and control monoclonal antibody (mAb) (Fig. 8). These results suggested that CD8-positive cells do not contribute to conferring anti-SFTSV protective cellular immunity.

**Fig 7.**
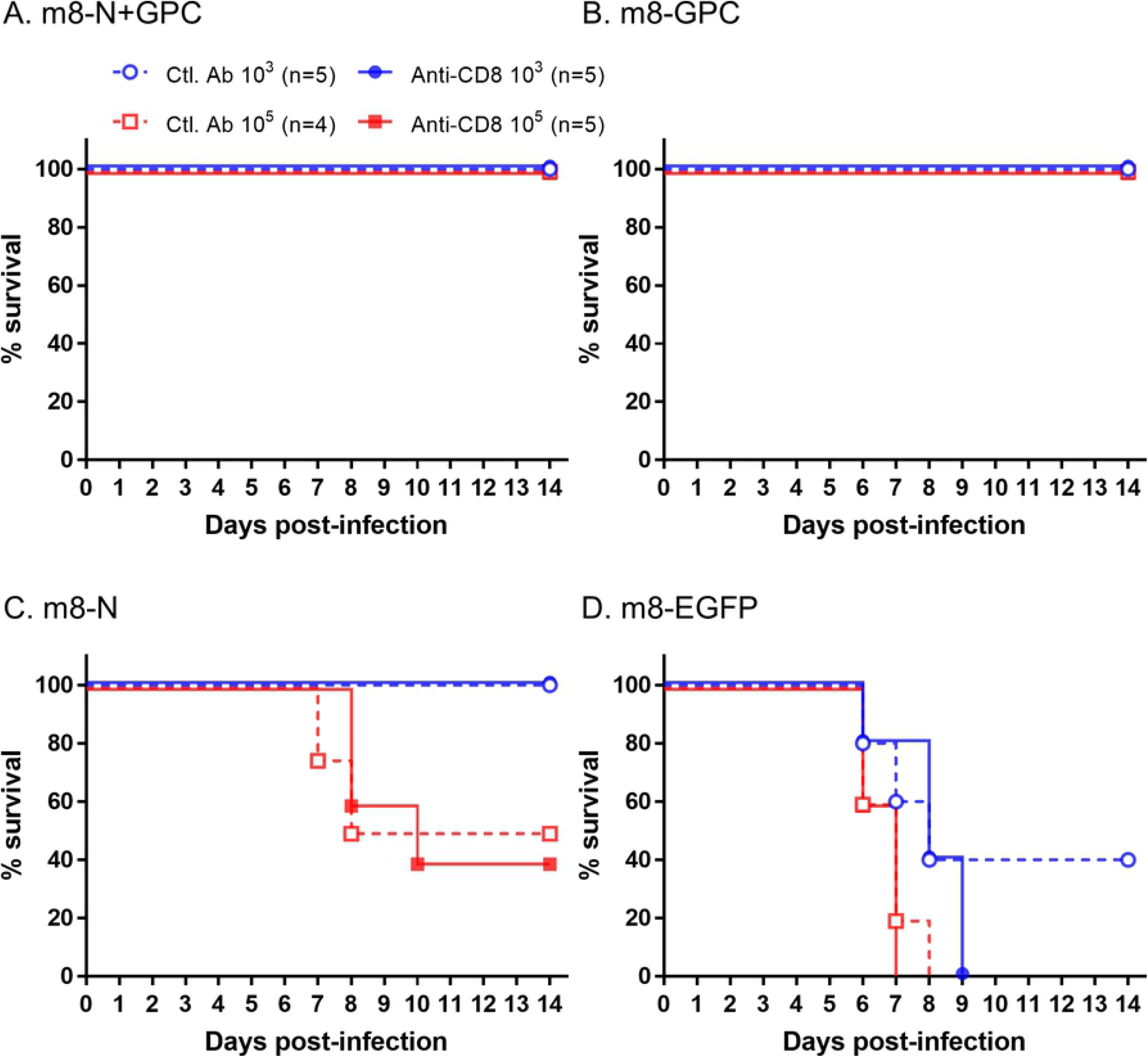
Survival in m8-SFTSV gene-inoculated mice following lethal SFTSV challenge with CD8-positive cell depletion. IFNAR-/- mice were subcutaneously inoculated twice at a 2-week interval with a dose of 1 × 10^6^ PFU of the m8-based SFTSV vaccines. Two weeks after the second inoculation of mice with m8-N+GPC (A), m8-GPC (B), m8-N (C) or m8-EGFP (D), the mice were challenged with 1 × 10^3^ (blue circle) or 1 × 10^5^ (red square) TCID_50_ of SFTSV YG-1 and were inoculated with anti-CD8 (strait line with filled symbol) mAb, to deplete the CD8-positive cells, or control (dotted line with open symbol) mAb on −1, 2, 5, and 8 DPI before and after SFTSV challenge. Survival was evaluated daily for 2 weeks. The number of mice in a group is shown as “n” in the legend.

**Fig 8.**
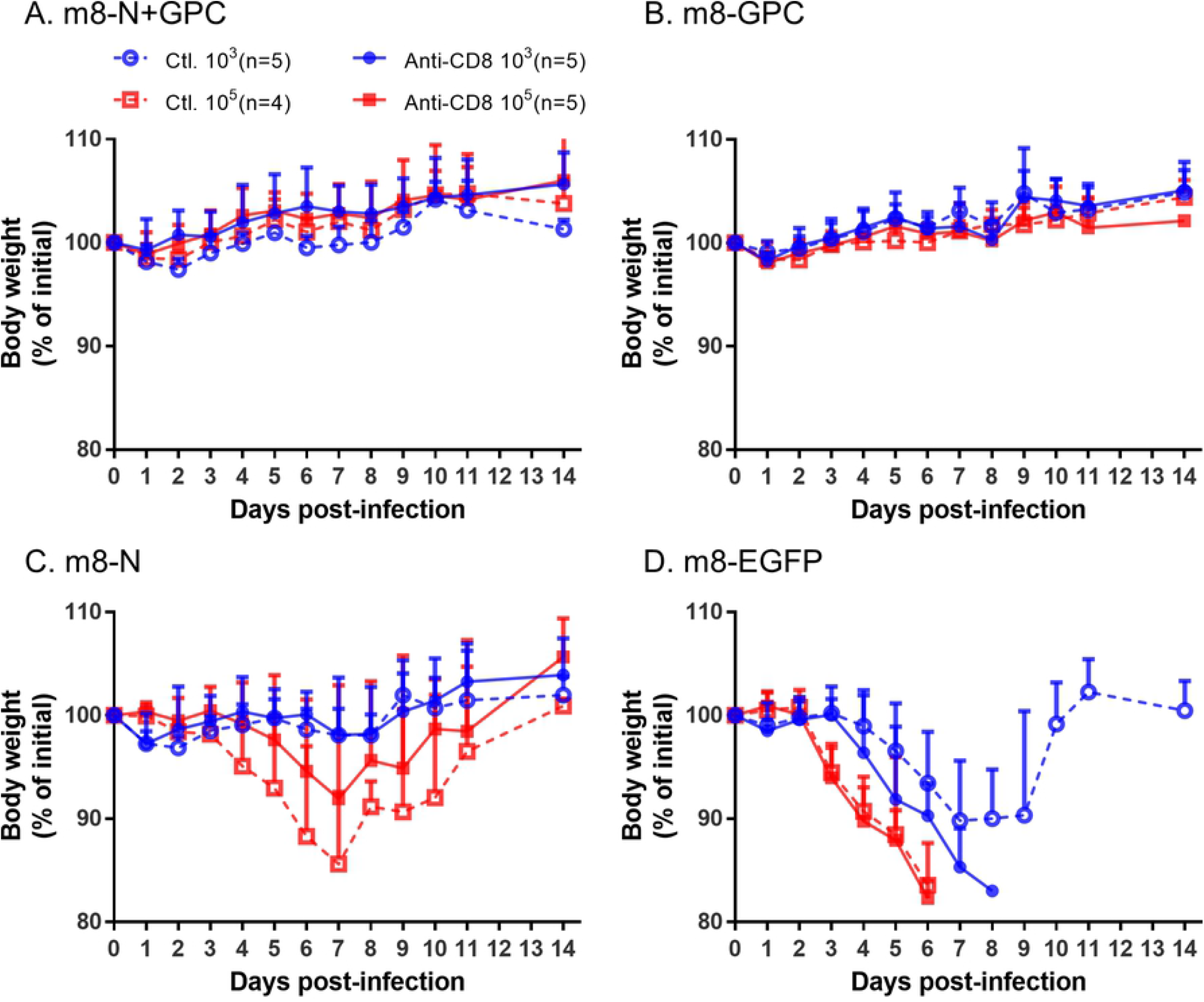
Weight change in m8-SFTSV gene-inoculated mice following a lethal SFTSV challenge with CD8-positive cell depletion. The inoculation schedule and figure legend are described in Fig. 7. The percent weight change from the initial weight was measured daily for 2 weeks.

## Discussion

An SFTS vaccine is urgently needed. One of the strategies for developing safe and effective vaccines is to use recombinant vaccine vectors. VAC has been served as a recombinant vaccine vector for many types of infectious diseases, including highly pathogenic infectious diseases [17]. In particular, MVA-based recombinant vaccines for CCHF and Ebola virus disease were reported to be effective in animal models [26–28]. Since there are many similarities between CCHF and SFTS (e.g., *Bunyavirales* order, tick-borne infectious disease, viral hemorrhagic fever, and other clinical findings) [29, 30], we hypothesized that an m8-based recombinant vaccine might be effective for SFTS.

Interestingly, this study elucidated both m8-N and m8-GPC conferred protection against lethal SFTSV challenge in mice, although m8-N appeared less effective than m8-GPC. Whereas CCHF virus (CCHFV) GPC-expressing MVA but not CCHF virus (CCHFV) N-expressing MVA conferred protection against lethal CCHFV challenge [26, 27]. It is hypothesized that the active acquired immunity elicited by SFTSV N and CCHFV N is dominantly cellular immunity, since the specific antibody cannot access the N in the virion. Therefore, cellular immunity against SFTSV infection could play a more crucial role than that of CCHFV infection.

The expression of the GPC gene in the cells infected with the Uukuniemi virus and Rift Valley fever virus in *Phenuiviridae* is necessary and sufficient for forming the VLP [31–33]. Since VLP presents antigens densely and repetitively in general, VLP is advantageous for vaccine use as it is efficiently taken up and processed by antigen-presenting cells, and also activates B cells via the physical crosslinking of B cell receptors [34, 35]. In this study, it was confirmed that the SFTSV VLP was produced in the cells *in vitro* when infected with m8-GPC or m8-N+GPC. As for the live recombinant SFTSV vaccines, m8-GPC and m8-N+GPC have the advantage of VLP production, which might occur not only *in vitro* but also *in vivo*, eliciting strong cellular and antibody responses. Although the immunogenicity (in terms of the humoral immunity) in mice induced by authentic SFTSV infection was slightly higher than that induced by recombinant m8-based SFTSV vaccines (Fig. 2), the immune response induced by vaccination with m8-GPC or m8-N+GPC completely protected against lethal SFTSV infection (Fig. 3). This result suggests that the mechanism through which m8-N+GPC induced protective immunity to SFTSV infection by mimicked that of authentic SFTSV infection.

The main target population for SFTS vaccination is people over 50 years of age, since age is a critical risk factor for suffering from severe and lethal SFTS [4, 36–38]. All countries terminated the routine smallpox vaccine program for infants by 1980. For example, the routine smallpox vaccination program was terminated in Japan in 1976. Since the youngest individuals who received the smallpox vaccine are now in their 40s, preexisting immunity to VAC is a considerable factor that might affect the ability of m8-based vaccines to induce protection against SFTSV. This study demonstrated the significant efficacy of the m8-GPC and m8-N+GPC vaccine in mice pre-vaccinated with VAC, even though there was only a one-month interval between the VAC (Lister) inoculation and m8-based SFTSV vaccines inoculation (Fig. 4). There have been no orthopoxvirus epidemics in the more than 40 years since people who received the smallpox vaccine received their last vaccination. Thus, it is highly likely that m8-based SFTSV vaccines will still be effective for individuals who have previously received the smallpox vaccine.

In general, recombinant VAC elicits both humoral (i.e., antibodies) and cellular immunity (i.e., cytotoxic T lymphocytes) against a target antigen [26, 27]. The key players in the humoral and cellular immunity are antibodies and cytotoxic CD8-positive T lymphocytes. In the present study, it was suggested that the specific antibodies against SFTSV GPC induced by m8-GPC or m8-N+GPC—but not those against N induced by m8-N—contributed to the protection against the SFTSV challenge in mice (Fig. 6). Hence, the major immunity that played a role in the protection of mice from the lethal SFTSV challenge induced by m8-N might be cellular immunity. Furthermore, cellular immunity induced by m8-GPC and m8-N+GPC may also play a role. However, SFTSV gene-specific CD8-positive cells elicited by m8-based SFTSV vaccines were dispensable for the protection against the SFTSV challenge. Further studies are needed to clarify the role of cellular immunity in the protection of subjects, including humans, from SFTSV infection.

Interestingly, it was previously reported that *in vivo* depletion of CD8-positive T lymphocytes in rats immunized with recombinant VAC that expresses measles genes did not alter the protective effect against a measles virus challenge [39, 40]. Although the authors did not test the *in vivo* depletion of CD4-positive cells, they hypothesized that the CD4-positive T cell-mediated immune response specific for measles gene products was sufficient to control measles virus infection. The contribution of CD4-positive T cells in mice inoculated with each of the m8-based SFTSV vaccines during the SFTSV challenge should be studied.

In summary, a novel recombinant m8 that expresses SFTSV N, GPC, or both N and GPC was generated. m8-GPC-infection and m8-N+GPC-infection of the cells induced the production of SFTSV-like-particles. In addition to m8-GPC and m8-N+GPC, m8-N was confirmed to confer protective efficacy against lethal SFTSV infection in mice. We concluded that m8-based SFTSV vaccines are promising candidates for an SFTS vaccine.

## Materials and methods

### Ethics statement

All experiments associated with animals were performed in animal biological safety level 2 or 3 containment laboratories at the National Institute of Infectious Diseases in Japan (NIID) under strict regulations of the animal experimentation guidelines of the NIID. The protocol was approved by the Institutional Animal Care and Use Committee of the NIID (no. 117059, 117130, 117131, 118127, 118129 and 118173).

### Cells and viruses

293FT cells (Thermo Fisher Scientific, Waltham, MA), Vero cells, and RK13 cells were grown in Dulbecco’s Modified Eagle’s Medium (DMEM; Wako, Osaka, Japan) supplemented with 5% of heat-inactivated fetal bovine serum (FBS) and antibiotics (Sigma-Aldrich Japan, Tokyo). VAC strain Lister, which is the grandparental strain of m8 (accession No. AY678276) [41], m8 (accession No. AY678275), and recombinant m8 expressing an enhanced green fluorescent protein (m8-EGFP), which was generated previously [42] were propagated in RK13 cells. The infectious titer in RK13 cells was determined by a standard plaque-forming unit (PFU) assay [41]. SFTSV strains YG-1 (accession No. AB817979, AB817987, AB817995), SPL005 (AB817982, AB817990, AB817998), and SPL010 (AB817983, AB817991, AB817999) were propagated in Vero cells. The infectious dose in Vero cells was determined with the 50% standard tissue culture infectious dose (TCID_50_) assay, with visualization of infection by an indirect immunofluorescence assay, as described previously [13]. Briefly, cells infected with SFTSV were reacted with an in-house-made rabbit anti-SFTSV N polyclonal antibody, then stained with Alexa Fluor 488-conjugated goat anti-rabbit IgG H+L antibody (Thermo Fisher Scientific).

### Plasmids

The open reading frames (ORFs) of N and GPC of SFTSV strain SPL005 were amplified using RNA purified from cells infected with SFTSV by two-step conventional RT-PCR. These ORFs containing a synthetic VAC early/late promoter [43] were also cloned into the NotI (N) or an XhoI (GPC) site of precB5R, a plasmid for generating recombinant m8s by homologous recombination [42]. The plasmids inserted with N and GPC genes were named precB5R-N and precB5R-GPC, respectively. The ORF of N with the vaccinia early/late promoter was inserted into the NotI site of precB5R-GPC to construct a plasmid named precB5R-N+GPC that contained both ORFs: N and GPC. The mammalian expression plasmid pHEK293 Ultra Expression Vector II (Takara Bio Inc., Shiga, Japan) was also inserted with the N or GPC ORF at the BamHI site of multicloning sites. Plasmids with the insertion of N and GPC were named pHEK293-N and -GPC, respectively. The N protein expression plasmid was inserted with an N gene with a FLAG-tag at the C-terminus of the gene.

### Generation of recombinant m8 harboring SFTSV genes

Recombinant m8, m8-N, m8-GPC and m8-N+GPC, with the insertion of only the N gene, only the GPC gene, and both the N and GPC genes, respectively, in the flanking region between B4R and B6R genes, were generated with a method employing homologous recombination for foreign gene insertion, as described previously [42]. Briefly, 293FT cells were transfected with each of the following plasmids: precB5R-N, precB5R-GPC, and precB5R-N+GPC by using X-tremeGENE 9 (Roche Diagnostics K.K., Tokyo, Japan). Cells transfected with each plasmid were then infected with m8 at a multiplicity of infection (MOI) of 0.05 per cell. The cells were cultured in DMEM supplemented with 5% FBS for 3 days. The culture medium, along with the cells, was then collected and was freeze-thawed three times to prepare crude intermediate recombinant m8 stocks. RK13 cell monolayers were inoculated with each of the crude intermediate recombinant m8 stocks. Then, the inoculated stock was removed, and the cells were overlaid with Eagle’s Minimal Essential Medium (EMEM, Wako) containing 2% of FBS, 20 μg/ml mycophenolic acid (MPA, Sigma-Aldrich), 250 μg/ml xanthine, 15 μg/ml hypoxanthine, and 1% agarose M.E. (Iwai Chemicals, Tokyo, Japan). After three days, the mCherry fluorescence-positive plaques were identified by either fluorescence microscopy or an LED transilluminator (GELmieru, WAKO) at the wavelength of 500 nm. One fluorescence-positive plaque in the agarose plug was collected and mixed with DMEM supplemented with 5% FBS to prepare stocks of the intermediate recombinant m8 clones. RK13 cells in a monolayer in a 96-well plate were inoculated with the intermediate clones to confirm the expression capability of SFTSV N and/or GPC. The cells inoculated were fixed with a 1:1 methanol: acetone mixture at two days post-infection (DPI). The cells were then reacted with either an in-house-made rabbit anti-SFTSV N polyclonal antibody or an in-house mouse anti-SFTSV Gc mAb clone C6C1, followed by reaction with DyLight 594-conjugated goat anti-rabbit IgG H+L antibody (Abcam, Cambridge, UK) and DyLight 488-conjugated goat anti-mouse IgG H+L antibody (Abcam), respectively. The fluorescence-positive expression clones were then further purified by plaque isolation at least three times. RK13 cells were infected with the purified intermediate clones without supplementation with MPA, xanthine, and hypoxanthine to prepare the crude final recombinant m8 stocks. Final recombinant m8-based SFTSV vaccines, from which the selection marker genes were self-excised, resulting in their deletion, were cloned by selecting fluorescence-negative plaques. Hence, monolayered RK13 cells were infected with one of the final crude recombinant m8 stocks, and the cells were overlaid with agarose containing MEM supplemented with 2% FBS without supplementation with MPA, xanthine, and hypoxanthine. After three days, the mCherry fluorescence-negative plaques were collected under an LED transilluminator. The N and/or GPC expression-positive and the mCherry fluorescence-negative clones were next purified by plaque isolation at least three times to establish the final recombinants. The m8-N, m8-GPC, m8-N+GPC, and m8-EGFP were propagated in RK13 cells, and their infectious doses were determined with the standard plaque assay.

### Confirmation of SFTS virus-like-particles (VLP)

RK13 cells were infected with m8-GPC or m8-N+GPC at an MOI of 0.1. The culture supernatant was collected at 3 DPI and mixed with polyethylene glycol (PEG) 6000 at a final concentration of 8 w/v % for pelleting the VLP, and the mixture was incubated at 4°C overnight. The PEG precipitates were pelleted by centrifugation for 30 min at 1,500 ×g at 4 °C in a swing-out rotor. The pellets were resuspended with DMEM supplemented with 5% FBS. The VLP was purified through a double-cushion of 20% and 60% sucrose in H_2_O in an SW-41 swing-out rotor by ultracentrifugation (Beckman Coulter, Brea, CA) for 2 hours at 25,000 rpm at 4 °C in a swing-out rotor. The accumulated VLP that was banded at the 20/60% interface was negatively stained with 2% phosphotungstic acid and then observed using a JEM-1400 transmission electron microscope (JEOL, Tokyo, Japan).

### Immunoprecipitation

The incorporation of SFTSV N into the VLP was confirmed by immunoprecipitation. Briefly, RK13 cells were infected with m8-EGFP, m8-N, or m8-N+GPC at an MOI of 0.05. The culture supernatants were collected at 3 DPI. The remaining cells were lysed with sodium dodecyl sulfate (SDS) sample buffer and heated at 95 °C for 5 min. One milliliter of each supernatant was incubated with or without 3 μg of mouse anti-SFTSV Gc mAb (clone C6C1) with rotation for 1 hour at room temperature followed by further incubation for 1 hour with the addition of 15 μl of goat anti-mouse IgG magnetic beads (New England Biolabs, Ipswich, MA, USA). The magnetic beads in the supernatant were collected on a magnetic stand and washed 2 times with PBS. Twenty microliters of SDS sample buffer was added to the beads and heated at 95 °C for 5 min. The infected cell lysates and immunoprecipitated samples were fractionated by SDS polyacrylamide gel electrophoresis (SDS-PAGE) and subjected to Western blotting. The membrane was reacted with rabbit anti-SFTSV N polyclonal antibody, followed by reaction with horseradish peroxidase (HRP)-conjugated goat anti-rabbit IgG (H+L) antibody (KPL, Gaithersburg, MD). The immunoreactive proteins were visualized by LAS-4000mini (Fujifilm, Tokyo, Japan) with ECL Prime Western blotting detection reagents (G.E. Healthcare, Chicago, IL, USA).

### Animals

IFNAR-/- mice on the C57BL/6 background were used for all of the animal experiments. The IFNAR-/- mice were previously produced by mating DNase II/type I interferon receptor double-knockout mice (strain B6.129-Dnase2a<tm1Osa> Ifnar1<tm1Agt>) [44–46] and C57BL/6 mice (age, 4–10 weeks; sex, male or female) and bred under specific pathogen-free (SPF) conditions at the NIID [47].

### Evaluation of vaccine efficacy in VAC-immunonegative mice

The mice were subcutaneously inoculated twice at a two-week interval with 1 × 10^6^ PFU of m8-EGFP, m8-N, m8-GPC, or m8-N+GPC in a volume of 100 μl per mouse under anesthesia with midazolam, medetomidine and butorphanol tartrate combination. At two weeks after the second inoculation, the mice were euthanized with Isoflurane and sacrificed to collect blood to evaluate immunogenicity or subcutaneously challenged with 1 × 10^3^ or 1 × 10^5^ TCID_50_ of SFTSV strain YG-1 in a volume of 100 μl per mouse under anesthesia with midazolam, medetomidine and butorphanol tartrate combination to evaluate the protective efficacy of each m8-based SFTSV vaccine candidate. The body weight and clinical signs of the mice challenged with SFTSV was monitored daily for two weeks. In the experiments challenged with SFTSV, the mice were euthanized with Isoflurane when the mice became moribund or expected to be die since of the difficulty in eating and/or drinking.

### Evaluation of vaccine efficacy in VAC-immunopositive mice

Four- to five-week-old IFNAR-/- mice (10 per group) were subcutaneously inoculated with 1 × 10^6^ PFU of a VAC strain Lister, a second-generation smallpox vaccine strain, in a volume of 100 μl per mouse at 4 weeks prior to the first inoculation with the m8-based SFTSV vaccine. The VAC pre-inoculated mice were then subcutaneously inoculated twice at a two-week interval with each m8-based SFTSV vaccine followed by subcutaneous challenge with 1 × 10^3^ or 1 × 10^5^ TCID_50_ of SFTSV YG-1, as described above. The body weight and clinical signs of the mice challenged with SFTSV was monitored daily for two weeks.

### Evaluation of the anti-SFTSV antibody transfer efficacy and cellular immunity in mice

The following experiment was designed to evaluate the humoral immunity induced by vaccination with m8-based SFTSV vaccines. Sera were collected from mice subcutaneously inoculated twice with 1 × 10^6^ PFU of m8-EGFP, m8-N, m8-GPC, or m8-N+GPC at a two-week interval, as described above. Naïve mice were passively transferred twice with 100 μl of sera at 1 day and immediately before the subcutaneous SFTSV challenge at 1 × 10^3^ TCID_50_. to evaluate the cellular immunity induced by based SFTSV vaccines, CD8-positive cells were depleted *in vivo* by the administration of anti-CD8 mAb, as described previously [48]. It was confirmed that 96.1 and 99.2% of CD8-positive cells were depleted from the spleen in naïve IFNAR-/- mice at 1 and 3 days post-mAb inoculation, respectively, when 250 μg of rat anti-mouse CD8 mAb clone 2.43 (BioXCell, West Lebanon, NH) intraperitoneally administered. Rat anti-keyhole limpet hemocyanin mAb clone LTF-2 (BioXCell) was used as the control antibody. The 6- to 9-week-old IFNAR-/- mice (4 to 5 per group) were subcutaneously inoculated twice with 1 × 10^6^ PFU of m8-EGFP, m8-N, m8-GPC, or m8-N+GPC at a 2-week interval, as described above. Two weeks later, the mice were subcutaneously challenged with 1 × 10^3^ or 1 × 10^5^ TCID_50_ of SFTSV YG-1 and were intraperitoneally inoculated with 250 μg of anti-CD8 mAb clone 2.43 on day −1, 2, 5, and 8 after SFTSV challenge, taking the day on which the mice were challenged with SFTSV as day 0. The body weight and clinical signs of the mice challenged with SFTSV was monitored daily for two weeks.

### Indirect immunofluorescence assay

IFA antigen-spotted slides, 293FT cells were transfected with either pHEK293-N or pHEK293-GPC with an expression enhancer plasmid, pHEK293 (Takara Bio Inc.). A mixture of the 293FT cells transfected with expression vectors in combination and untransfected 293FT cells at a ratio of 1:3 were washed with PBS, spotted on glass slides (Matsunami glass IND., Ltd., Osaka, Japan), dry-fixed, and treated with acetone. To measure the titer of the SFTSV N- or GPC-specific IgG in mice, serum samples immobilized at 56 °C for 30 min were serially diluted two-fold and added onto the antigens on the glass slides. The antigens were then reacted with an Alexa Fluor 488-conjugated goat anti-mouse IgG H+L antibody (Thermo Fisher Scientific). The antibody titer was defined as the reciprocal of the highest dilution level at which a specific fluorescent signal was detected.

### Focus reduction neutralization test

To measure the titer of neutralizing antibody to SFTSV in mouse serum, 100 TCID_50_ aliquots of SFTSV were mixed with 4-fold serially diluted immobilized sera, and the mixtures were incubated for 1 hour. Confluent Vero cell cultures in 12-well microtiter plates were inoculated with each virus and serum mixture for 1 hour, overlaid with DMEM containing 2% FBS and 1% methylcellulose, and then cultured for 4 to 5 days. The cells were fixed with PBS containing 10% formalin and were reacted with rabbit anti-SFTSV N polyclonal antibody. The foci of SFTSV infection were then visualized using a standard immunoperoxidase method. The percent N.T. antibody titer was determined to divide the number of foci in the well inoculated with the virus and serum from m8-SFTSV gene or SFTSV inoculated mouse by that of naïve mice.

### Quantitative one-step RT-PCR

Specific quantitative one-step reverse transcription-PCR (qPCR) was performed as reported previously [49]. Specific qPCR primer and probe sets targeted to the N gene (forward; TGTCAGAGTGGTCCAGGATT, reverse; ACCTGTCTCCTTCAGCTTCT, probe; FAM-TGGAGTTTGGTGAGCAGCAGC-BHQ1) were used. Total RNAs were extracted from 200 μl of mice sera using a High Pure viral RNA kit (Roche Applied Science) according to the manufacturer’s protocol. The elution volume for RNA extraction was 50 μl. For the qPCR assay, an aliquot of the extracted RNA solution was added to the reaction mixture for the QuantiTect probe RT-PCR kit (Qiagen, Hilden, Germany), which contained 2x QuantiTect probe RT-PCR master mix, QuantiTect RT mix, H2O, and 10x primer-probe mix, which contained 4 μM of each specific primer, 2 μM TaqMan probe(s), and 2 μl of contamplicon probe. After PCR activation at 95°C for 15 min, the reverse transcription reaction was carried out at 50°C for 30 min, followed by 45 cycles of amplification under the following conditions: 94°C for 15 s and 60°C for 60 s in a LightCycler 96 (Roche).

### Histopathology and immunohistochemistry

As described above, IFNAR-/- mice were subcutaneously inoculated with 1 × 10^6^ PFU of the m8-EGFP or each m8-based SFTSV vaccine twice at a 2-week interval and were intraperitoneally challenged with 1 × 10^5^ TCID_50_ of SFTSV YG-1, 2 weeks after the second inoculation. The SFTSV-infected mice were euthanized with Isoflurane and sacrificed at 1, 3, and 5 DPI, and the blood, cervical lymph nodes, spleen, liver, and kidneys were collected. The tissues were fixed with PBS containing 10% formalin and embedded with paraffin to prepare blocks. Sections were prepared from the tissue block and were stained with hematoxylin-eosin for a histopathological examination. An immunohistochemical analysis to detect SFTSV N protein was performed as previously described [13]. The sections were first reacted with a rabbit anti-SFTSV N polyclonal antibody. Antigens were retrieved by hydrolytic autoclaving in citrate buffer (pH 6.0) for 10 min at 121°C. IHC staining was then performed using a standard immunoperoxidase method.

### Statistical analysis

All statistical analyses were performed using GraphPad Prism 7 (GraphPad Software, La Jolla, CA). Survival curves were plotted according to a Kaplan-Meier analysis, and the protective efficacy of the m8-based SFTSV vaccines against SFTSV challenge in mice was evaluated using a log-rank test with Bonferroni correction to adjust the significance level for multiple comparisons. The equality of the means of the virulent virus titer or the viral RNA levels in mouse sera was evaluated using a two-way analysis of variance (ANOVA) with Sidak’s multiple-comparison test. P values of <0.05 were considered to indicate statistical significance.

## Acknowledgments

We would like to thank Mr. Ken-Ichi Shibasaki, Ms. Momoko Ogata, Ms. Yoshiko Fukui and Ms. Mihoko Tsuda for their excellent technical assistance.

## References

1. Yu XJ, Liang MF, Zhang SY, Liu Y, Li JD, Sun YL, et al. Fever with thrombocytopenia associated with a novel bunyavirus in China. N Engl J Med. 2011;364(16):1523–32. Epub 2011/03/18. doi: 10.1056/NEJMoa1010095. PubMed PMID: 21410387; PubMed Central PMCID: PMC3113718.

2. Xu B, Liu L, Huang X, Ma H, Zhang Y, Du Y, et al. Metagenomic analysis of fever, thrombocytopenia and leukopenia syndrome (FTLS) in Henan Province, China: discovery of a new bunyavirus. PLoS Pathog. 2011;7(11):e1002369. Epub 2011/11/25. doi: 10.1371/journal.ppat.1002369. PubMed PMID: 22114553; PubMed Central PMCID: PMC3219706.

3. Zhang YZ, Zhou DJ, Xiong Y, Chen XP, He YW, Sun Q, et al. Hemorrhagic fever caused by a novel tick-borne Bunyavirus in Huaiyangshan, China. Zhonghua Liu Xing Bing Xue Za Zhi. 2011;32(3):209–20. Epub 2011/04/05. PubMed PMID: 21457654.

4. Kato H, Yamagishi T, Shimada T, Matsui T, Shimojima M, Saijo M, et al. Epidemiological and Clinical Features of Severe Fever with Thrombocytopenia Syndrome in Japan, 2013-2014. PLoS One. 2016;11(10):e0165207. Epub 2016/10/25. doi: 10.1371/journal.pone.0165207. PubMed PMID: 27776187; PubMed Central PMCID: PMCPMC5077122.

5. Reece LM, Beasley DW, Milligan GN, Sarathy VV, Barrett AD. Current status of Severe Fever with Thrombocytopenia Syndrome vaccine development. Curr Opin Virol. 2018;29:72–8. Epub 2018/04/12. doi: 10.1016/j.coviro.2018.03.005. PubMed PMID: 29642053.

6. Choi SJ, Park SW, Bae IG, Kim SH, Ryu SY, Kim HA, et al. Severe Fever with Thrombocytopenia Syndrome in South Korea, 2013-2015. PLoS neglected tropical diseases. 2016;10(12):e0005264. doi: 10.1371/journal.pntd.0005264. PubMed PMID: 28033338; PubMed Central PMCID: PMCPMC5226827.

7. Hu J, Li S, Zhang X, Zhao H, Yang M, Xu L, et al. Correlations between clinical features and death in patients with severe fever with thrombocytopenia syndrome. Medicine (Baltimore). 2018;97(22):e10848. Epub 2018/06/01. doi: 10.1097/MD.0000000000010848. PubMed PMID: 29851797; PubMed Central PMCID: PMCPMC6392624.

8. Shin J, Kwon D, Youn SK, Park JH. Characteristics and Factors Associated with Death among Patients Hospitalized for Severe Fever with Thrombocytopenia Syndrome, South Korea, 2013. Emerg Infect Dis. 2015;21(10):1704–10. Epub 2015/09/25. doi: 10.3201/eid2110.141928. PubMed PMID: 26402575; PubMed Central PMCID: PMCPMC4593431.

9. Ding YP, Liang MF, Ye JB, Liu QH, Xiong CH, Long B, et al. Prognostic value of clinical and immunological markers in acute phase of SFTS virus infection. Clin Microbiol Infect. 2014;20(11):O870–8. Epub 2014/04/02. doi: 10.1111/1469-0691.12636. PubMed PMID: 24684627.

10. Liu S, Chai C, Wang C, Amer S, Lv H, He H, et al. Systematic review of severe fever with thrombocytopenia syndrome: virology, epidemiology, and clinical characteristics. Reviews in medical virology. 2014;24(2):90–102. doi: 10.1002/rmv.1776. PubMed PMID: 24310908; PubMed Central PMCID: PMC4237196.

11. Deng B, Zhang S, Geng Y, Zhang Y, Wang Y, Yao W, et al. Cytokine and chemokine levels in patients with severe fever with thrombocytopenia syndrome virus. PLoS One. 2012;7(7):e41365. Epub 2012/08/23. doi: 10.1371/journal.pone.0041365. PubMed PMID: 22911786; PubMed Central PMCID: PMCPMC3404083.

12. Kim KH, Yi J, Kim G, Choi SJ, Jun KI, Kim NH, et al. Severe fever with thrombocytopenia syndrome, South Korea, 2012. Emerg Infect Dis. 2013;19(11):1892–4. doi: 10.3201/eid1911.130792. PubMed PMID: 24206586; PubMed Central PMCID: PMC3837670.

13. Takahashi T, Maeda K, Suzuki T, Ishido A, Shigeoka T, Tominaga T, et al. The first identification and retrospective study of Severe Fever with Thrombocytopenia Syndrome in Japan. J Infect Dis. 2014;209(6):816–27. doi: 10.1093/infdis/jit603. PubMed PMID: 24231186.

14. Tran XC, Yun Y, Van An L, Kim SH, Thao NTP, Man PKC, et al. Endemic Severe Fever with Thrombocytopenia Syndrome, Vietnam. Emerg Infect Dis. 2019;25(5):1029–31. Epub 2019/04/20. doi: 10.3201/eid2505.181463. PubMed PMID: 31002059; PubMed Central PMCID: PMCPMC6478219.

15. Dong F, Li D, Wen D, Li S, Zhao C, Qi Y, et al. Single dose of a rVSV-based vaccine elicits complete protection against severe fever with thrombocytopenia syndrome virus. NPJ Vaccines. 2019;4:5. Epub 2019/02/01. doi: 10.1038/s41541-018-0096-y. PubMed PMID: 30701094; PubMed Central PMCID: PMCPMC6347601 All the remaining authors declare no competing interests.

16. Kwak JE, Kim YI, Park SJ, Yu MA, Kwon HI, Eo S, et al. Development of a SFTSV DNA vaccine that confers complete protection against lethal infection in ferrets. Nat Commun. 2019;10(1):3836. Epub 2019/08/25. doi: 10.1038/s41467-019-11815-4. PubMed PMID: 31444366; PubMed Central PMCID: PMCPMC6707330.

17. Walsh SR, Dolin R. Vaccinia viruses: vaccines against smallpox and vectors against infectious diseases and tumors. Expert Rev Vaccines. 2011;10(8):1221–40. doi: 10.1586/erv.11.79. PubMed PMID: 21854314; PubMed Central PMCID: PMCPMC3223417.

18. Ramezanpour B, Haan I, Osterhaus A, Claassen E. Vector-based genetically modified vaccines: Exploiting Jenner’s legacy. Vaccine. 2016;34(50):6436–48. Epub 2016/12/29. doi: 10.1016/j.vaccine.2016.06.059. PubMed PMID: 28029542; PubMed Central PMCID: PMCPMC7115478.

19. Lane JM, Ruben FL, Neff JM, Millar JD. Complications of smallpox vaccination, 1968: results of ten statewide surveys. J Infect Dis. 1970;122(4):303–9. Epub 1970/10/01. doi: 10.1093/infdis/122.4.303. PubMed PMID: 4396189.

20. Kenner J, Cameron F, Empig C, Jobes DV, Gurwith M. LC16m8: an attenuated smallpox vaccine. Vaccine. 2006;24(47-48):7009–22. doi: 10.1016/j.vaccine.2006.03.087. PubMed PMID: 17052815.

21. Belyakov IM, Earl P, Dzutsev A, Kuznetsov VA, Lemon M, Wyatt LS, et al. Shared modes of protection against poxvirus infection by attenuated and conventional smallpox vaccine viruses. Proc Natl Acad Sci U S A. 2003;100(16):9458–63. doi: 10.1073/pnas.1233578100. PubMed PMID: 12869693; PubMed Central PMCID: PMCPMC170940.

22. Earl PL, Americo JL, Wyatt LS, Eller LA, Whitbeck JC, Cohen GH, et al. Immunogenicity of a highly attenuated MVA smallpox vaccine and protection against monkeypox. Nature. 2004;428(6979):182–5. doi: 10.1038/nature02331. PubMed PMID: 15014500.

23. Wyatt LS, Earl PL, Eller LA, Moss B. Highly attenuated smallpox vaccine protects mice with and without immune deficiencies against pathogenic vaccinia virus challenge. Proc Natl Acad Sci U S A. 2004;101(13):4590–5. doi: 10.1073/pnas.0401165101. PubMed PMID: 15070762; PubMed Central PMCID: PMCPMC384791.

24. Kennedy JS, Gurwith M, Dekker CL, Frey SE, Edwards KM, Kenner J, et al. Safety and immunogenicity of LC16m8, an attenuated smallpox vaccine in vaccinia-naive adults. J Infect Dis. 2011;204(9):1395–402. Epub 2011/09/17. doi: 10.1093/infdis/jir527. PubMed PMID: 21921208; PubMed Central PMCID: PMCPMC3218648.

25. Saito T, Fujii T, Kanatani Y, Saijo M, Morikawa S, Yokote H, et al. Clinical and immunological response to attenuated tissue-cultured smallpox vaccine LC16m8. JAMA. 2009;301(10):1025–33. Epub 2009/03/13. doi: 10.1001/jama.2009.289. PubMed PMID: 19278946.

26. Dowall SD, Buttigieg KR, Findlay-Wilson SJ, Rayner E, Pearson G, Miloszewska A, et al. A Crimean-Congo hemorrhagic fever (CCHF) viral vaccine expressing nucleoprotein is immunogenic but fails to confer protection against lethal disease. Human vaccines & immunotherapeutics. 2016;12(2):519–27. Epub 2015/08/27. doi: 10.1080/21645515.2015.1078045. PubMed PMID: 26309231; PubMed Central PMCID: PMCPMC5049717.

27. Buttigieg KR, Dowall SD, Findlay-Wilson S, Miloszewska A, Rayner E, Hewson R, et al. A novel vaccine against Crimean-Congo Haemorrhagic Fever protects 100% of animals against lethal challenge in a mouse model. PLoS One. 2014;9(3):e91516. Epub 2014/03/14. doi: 10.1371/journal.pone.0091516. PubMed PMID: 24621656; PubMed Central PMCID: PMCPMC3951450.

28. Domi A, Feldmann F, Basu R, McCurley N, Shifflett K, Emanuel J, et al. A Single Dose of Modified Vaccinia Ankara expressing Ebola Virus Like Particles Protects Nonhuman Primates from Lethal Ebola Virus Challenge. Sci Rep. 2018;8(1):864. Epub 2018/01/18. doi: 10.1038/s41598-017-19041-y. PubMed PMID: 29339750; PubMed Central PMCID: PMCPMC5770434.

29. Shayan S, Bokaean M, Shahrivar MR, Chinikar S. Crimean-Congo Hemorrhagic Fever. Lab Med. 2015;46(3):180–9. Epub 2015/07/23. doi: 10.1309/LMN1P2FRZ7BKZSCO. PubMed PMID: 26199256.

30. Miyamoto S, Ito T, Terada S, Eguchi T, Furubeppu H, Kawamura H, et al. Fulminant myocarditis associated with severe fever with thrombocytopenia syndrome: a case report. BMC Infect Dis. 2019;19(1):266. Epub 2019/03/20. doi: 10.1186/s12879-019-3904-8. PubMed PMID: 30885147; PubMed Central PMCID: PMCPMC6423866.

31. Strandin T, Hepojoki J, Vaheri A. Cytoplasmic tails of bunyavirus Gn glycoproteins-Could they act as matrix protein surrogates? Virology. 2013;437(2):73–80. Epub 2013/01/30. doi: 10.1016/j.virol.2013.01.001. PubMed PMID: 23357734.

32. Piper ME, Sorenson DR, Gerrard SR. Efficient cellular release of Rift Valley fever virus requires genomic RNA. PLoS One. 2011;6(3):e18070. Epub 2011/03/30. doi: 10.1371/journal.pone.0018070. PubMed PMID: 21445316; PubMed Central PMCID: PMCPMC3061922.

33. Overby AK, Popov V, Neve EP, Pettersson RF. Generation and analysis of infectious virus-like particles of uukuniemi virus (bunyaviridae): a useful system for studying bunyaviral packaging and budding. J Virol. 2006;80(21):10428–35. Epub 2006/08/25. doi: 10.1128/JVI.01362-06. PubMed PMID: 16928751; PubMed Central PMCID: PMCPMC1641803.

34. Zabel F, Mohanan D, Bessa J, Link A, Fettelschoss A, Saudan P, et al. Viral particles drive rapid differentiation of memory B cells into secondary plasma cells producing increased levels of antibodies. J Immunol. 2014;192(12):5499–508. Epub 2014/05/14. doi: 10.4049/jimmunol.1400065. PubMed PMID: 24821969.

35. Li C, Liu F, Liang M, Zhang Q, Wang X, Wang T, et al. Hantavirus-like particles generated in CHO cells induce specific immune responses in C57BL/6 mice. Vaccine. 2010;28(26):4294–300. Epub 2010/05/04. doi: 10.1016/j.vaccine.2010.04.025. PubMed PMID: 20433802.

36. Fu Y, Li S, Zhang Z, Man S, Li X, Zhang W, et al. Phylogeographic analysis of severe fever with thrombocytopenia syndrome virus from Zhoushan Islands, China: implication for transmission across the ocean. Sci Rep. 2016;6:19563. doi: 10.1038/srep19563. PubMed PMID: 26806841; PubMed Central PMCID: PMCPMC4726339.

37. Ding S, Niu G, Xu X, Li J, Zhang X, Yin H, et al. Age is a critical risk factor for severe fever with thrombocytopenia syndrome. PLoS One. 2014;9(11):e111736. Epub 2014/11/05. doi: 10.1371/journal.pone.0111736. PubMed PMID: 25369237; PubMed Central PMCID: PMCPMC4219771.

38. Guo CT, Lu QB, Ding SJ, Hu CY, Hu JG, Wo Y, et al. Epidemiological and clinical characteristics of severe fever with thrombocytopenia syndrome (SFTS) in China: an integrated data analysis. Epidemiol Infect. 2016;144(6):1345–54. Epub 2015/11/07. doi: 10.1017/S0950268815002678. PubMed PMID: 26542444.

39. Brinckmann UG, Bankamp B, Reich A, ter Meulen V, Liebert UG. Efficacy of individual measles virus structural proteins in the protection of rats from measles encephalitis. J Gen Virol. 1991;72 (Pt 10):2491–500. Epub 1991/10/01. doi: 10.1099/0022-1317-72-10-2491. PubMed PMID: 1833505.

40. Bankamp B, Brinckmann UG, Reich A, Niewiesk S, ter Meulen V, Liebert UG. Measles virus nucleocapsid protein protects rats from encephalitis. J Virol. 1991;65(4):1695–700. Epub 1991/04/01. PubMed PMID: 1825854; PubMed Central PMCID: PMCPMC239973.

41. Morikawa S, Sakiyama T, Hasegawa H, Saijo M, Maeda A, Kurane I, et al. An attenuated LC16m8 smallpox vaccine: analysis of full-genome sequence and induction of immune protection. J Virol. 2005;79(18):11873–91. doi: 10.1128/JVI.79.18.11873-11891.2005. PubMed PMID: 16140764; PubMed Central PMCID: PMCPMC1212643.

42. Omura N, Yoshikawa T, Fujii H, Shibamura M, Inagaki T, Kato H, et al. A Novel System for Constructing a Recombinant Highly-Attenuated Vaccinia Virus Strain (LC16m8) Expressing Foreign Genes and Its Application for the Generation of LC16m8-Based Vaccines against Herpes Simplex Virus 2. Japanese journal of infectious diseases. 2018;71(3):229–33. doi: 10.7883/yoken.JJID.2017.458. PubMed PMID: 29709968.

43. Chakrabarti S, Sisler JR, Moss B. Compact, synthetic, vaccinia virus early/late promoter for protein expression. BioTechniques. 1997;23(6):1094–7. PubMed PMID: 9421642.

44. Yoshida H, Okabe Y, Kawane K, Fukuyama H, Nagata S. Lethal anemia caused by interferon-beta produced in mouse embryos carrying undigested DNA. Nat Immunol. 2005;6(1):49–56. Epub 2004/11/30. doi: 10.1038/ni1146. PubMed PMID: 15568025.

45. Kawane K, Ohtani M, Miwa K, Kizawa T, Kanbara Y, Yoshioka Y, et al. Chronic polyarthritis caused by mammalian DNA that escapes from degradation in macrophages. Nature. 2006;443(7114):998–1002. Epub 2006/10/27. doi: 10.1038/nature05245. PubMed PMID: 17066036.

46. Muller U, Steinhoff U, Reis LF, Hemmi S, Pavlovic J, Zinkernagel RM, et al. Functional role of type I and type II interferons in antiviral defense. Science. 1994;264(5167):1918–21. Epub 1994/06/24. doi: 10.1126/science.8009221. PubMed PMID: 8009221.

47. Tani H, Fukuma A, Fukushi S, Taniguchi S, Yoshikawa T, Iwata-Yoshikawa N, et al. Efficacy of T-705 (Favipiravir) in the Treatment of Infections with Lethal Severe Fever with Thrombocytopenia Syndrome Virus. mSphere. 2016;1(1). doi: 10.1128/mSphere.00061-15. PubMed PMID: 27303697; PubMed Central PMCID: PMCPMC4863605.

48. Jing W, Gershan JA, Johnson BD. Depletion of CD4 T cells enhances immunotherapy for neuroblastoma after syngeneic HSCT but compromises development of antitumor immune memory. Blood. 2009;113(18):4449–57. Epub 2009/02/03. doi: 10.1182/blood-2008-11-190827. PubMed PMID: 19182203; PubMed Central PMCID: PMCPMC2676098.

49. Yoshikawa T, Fukushi S, Tani H, Fukuma A, Taniguchi S, Toda S, et al. Sensitive and specific PCR systems for detection of both Chinese and Japanese severe fever with thrombocytopenia syndrome virus strains and prediction of patient survival based on viral load. Journal of clinical microbiology. 2014;52(9):3325–33. doi: 10.1128/JCM.00742-14. PubMed PMID: 24989600; PubMed Central PMCID: PMCPMC4313158.

